# Sex chromosome trisomies are not associated with atypical lateralisation for language

**DOI:** 10.1101/248286

**Authors:** Alexander C. Wilson, Dorothy V. M. Bishop

## Abstract

**Aim:** Individuals with a sex chromosome trisomy (SCT) have disproportionate problems with language compared to nonverbal skills. This may result from disruption to the typical left hemisphere bias for language processing. We tested the hypothesis that SCTs would be associated with reduced left lateralisation for language.

**Method:** In a cross-sectional design, language laterality was measured during an animation description task using functional transcranial Doppler ultrasonography (fTCD). Data were available for 75 children with an SCT (26 47,XXX girls, 25 47,XXY boys, and 24 47,XYY boys; mean age was 11;4 years, SD = 3;10 years), and 132 comparison children with typical karyotypes (69 boys; mean age was 9;1 years, SD = 1;7 years).

**Results:** Lateralisation for language did not differ between the SCT and comparison groups, either in mean laterality index or relative frequency of each laterality category (left, bilateral and right). There were no differences when splitting the SCT group by trisomy. Handedness also showed no group effects.

**Interpretation:** Our data provide no evidence for disrupted lateralisation for language in SCTs. The brain basis of the cognitive phenotype in SCTs is unlikely to be a failure of the left hemisphere to specialise for language, as previously suggested.

**What this paper adds:**

- Children with a sex chromosome trisomy (SCT) have typically lateralised language.
- This disconfirms theories linking language problems to hemispheric specialization in SCTs

---

Sex chromosome trisomies (SCTs) are common genetic anomalies where an individual has an extra X or Y chromosome alongside the typical 46 chromosomes: males with an SCT are 47,XXY or 47,XYY, and females are 47,XXX. The risk of neurodevelopmental problems, especially language impairments, is increased among people with an SCT, as noted by a systematic review^1^. Average verbal IQ is around 1 SD lower than controls, and speech therapy is often required for delayed language development. In boys with an extra X or Y, nonverbal ability is relatively unimpaired, whereas it is depressed in girls. With this pattern of greater verbal than nonverbal difficulties, SCTs may offer insights into Developmental Language Disorder, in particular through understanding the neurobiological mechanism that mediates between the genetic anomaly and the language problems. The leading hypothesis is that an extra sex chromosome affects the dosage of genes involved in the typical asymmetric development of the brain^2,3^, which may disrupt the usual left-sided bias for language function. A right-sided or bilateral pattern may develop instead, possibly predisposing the individual to language disorder.

The existing evidence for atypical lateralisation in individuals with an SCT is suggestive. We are aware of two studies that examined functional measures of activity in cerebral cortex, and found a reduced left bias at the group level^4,5^. A SPECT resting-state study showed more bilateral perfusion of parietal and temporal regions in 47,XXY men (n=9) compared to controls who showed a left bias^4^. In an fMRI study, mean laterality indices (LIs) averaging activation across three language tasks indicated a reduced left bias across language-relevant regions in 15 47,XXY individuals compared to 14 control men^5^. However, the dataset contained an outlier: one 47,XXY participant was strongly right-lateralised, whereas all other participants were left-lateralised. It is unclear whether the results were robust to this outlier. A further paper reported preliminary fMRI analysis of an 47,XXY sample (n=8) showing reduced left-sided activation during a language task^6^.

There are mixed findings regarding structural asymmetries. Reduced left temporal lobe volume has been found in two samples of 47,XXY individuals (n=15 though only after excluding left-handers^7^; n=10^8^). However, neither study conducted direct between-hemispheres comparisons. Two studies report no differences in asymmetry, despite reduced overall brain volume in those with an extra X chromosome, in a sample including each SCT (n=34)^9^ and a 47,XXY sample (n=65)^10^. DeLisi and colleagues found similar results, with bilateral reductions in frontal and temporal volume, but typical laterality in 11 47,XXY individuals, although three of four locations with reduced white matter integrity were left-sided (left internal capsule, left arcuate and bilateral anterior cingulate)^11^. Cerebral torque (the typical pattern of greater anterior volume in the right hemisphere and greater posterior volume in the left) has been examined in 47,XXY individuals (n=10), but did not differ significantly from controls^12^. However, this study found reduced rightward asymmetry in white matter connecting the anterior commissure to the frontal lobe, and a non-significant trend (*p* = .056) for reduced frontal asymmetry, in the 47,XXY group. Reduced rightward asymmetry in the medial occipital lobe and superior temporal gyrus has been found in 47,XXY men compared to control men (but not women)^13^.

Though most neuroimaging work has focussed on 47,XXY groups, a few further studies have considered other SCTs. Right hemispheric regional differences have been found in 47,XYY boys (n=10), even when controlling for greater overall volume in this group^14^. The right occipital lobe showed increased gray matter volume, while the insula, inferior frontal gyrus and superior temporal cortex volumes were decreased. Note, however, that direct comparisons between the hemispheres were not reported. A group of 47,XXX girls (n=35) had decreased overall volume and regional reductions in frontal and temporal cortices, though laterality was not significantly different to controls^15^. A final study examined whether a supernumerary sex chromosome affected cortical thickness in five sex chromosome aneuploidies (n=137), but found relatively typical torque, with only small foci of difference^16^.

While neuro-anatomical differences have been found in individuals with an SCT, structural asymmetries seem unaffected. By contrast, functional neuroimaging studies, as well as behavioural studies using dichotic listening^17,18^, provide some evidence of a reduced bias for left-sided language processing in 47,XXY individuals. This suggests that failure to establish a specialised language network in the left hemisphere may account for the verbal impairments of individuals with an extra sex chromosome^2,12^. However, evidence is limited to small exploratory studies, and calls for confirmation in a large sample. In this study, we predicted that children with an SCT would show a reduced bias for left-sided blood flow during a language task, compared to children with a typical karyotype. We also tested for reduced right-handedness in the SCT group, given the link between manual and language laterality.

## Methods

### Participants

Using a cross-sectional design, we compared language laterality and handedness in children with an SCT aged between 6;0 and 15;11 (n=75) with a group of twin children aged between 6;0 and 11;11 (n=132). Our previous paper gives details of the twin sample, which did not vary in laterality from singleborn children^19^. Since independence of observations cannot be assumed among twins, only one twin from each pair was included in the present analysis: the child arbitrarily labelled ‘twin 1’.

Out of 143 children with an SCT recruited into the study, we collected useable fTCD data from 75, as shown in Figure 1. Recruitment was via NHS Clinical Genetics centres, two support groups (Unique: the Rare Chromosome Support Group, and the Klinefelter Syndrome Association) and self-referral through social media. An inclusion criterion was that children were aware of their trisomy status. Some children had been diagnosed following postnatal testing motivated by neurodevelopmental/behavioural problems. The phenotype of these children may be more severe, potentially biasing the sample, and so we grouped them as a high-risk-of-bias subgroup. All other children formed the low-risk-of-bias subgroup.

**Figure 1.**
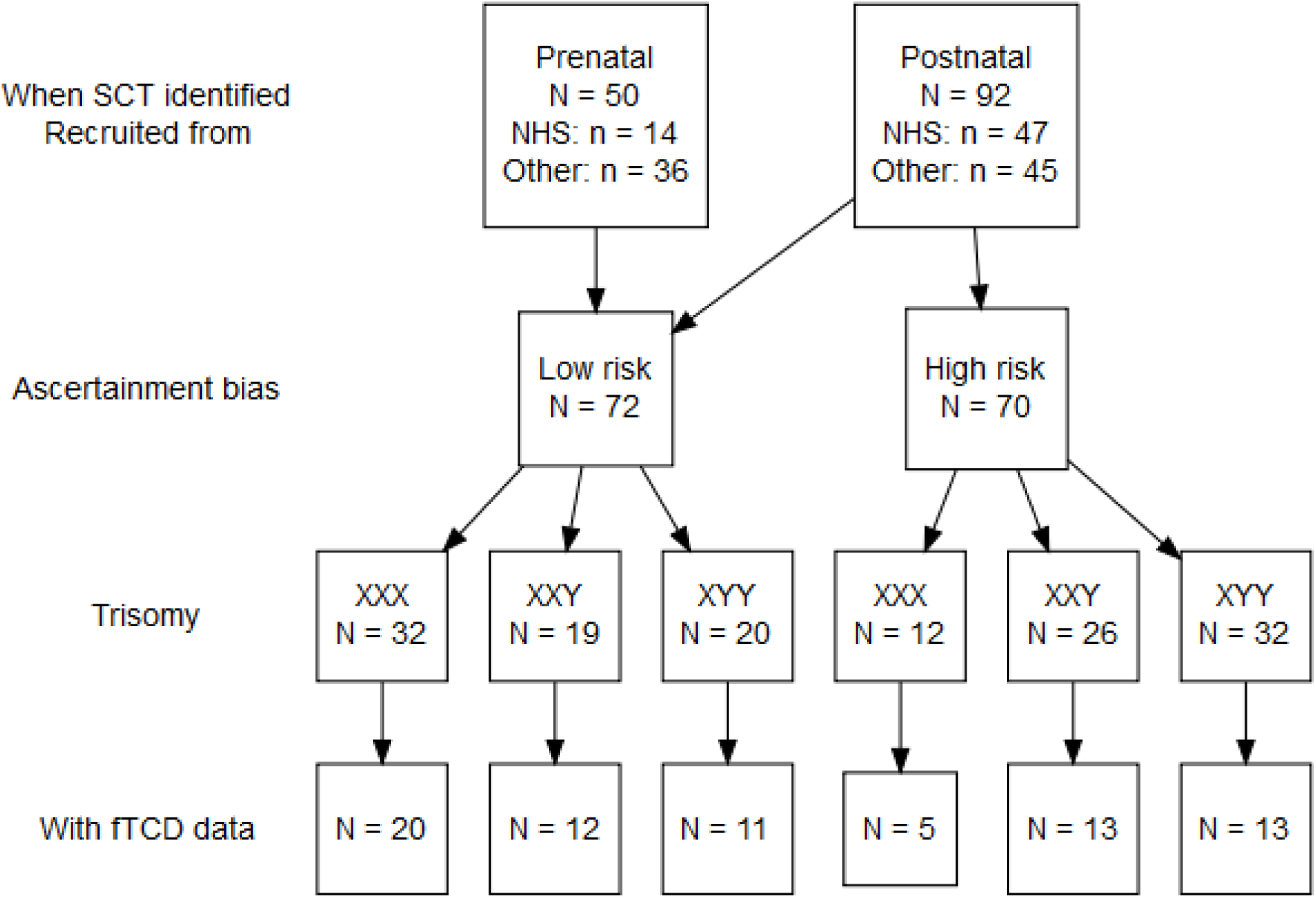
Chart showing the flow of children with an SCT through the study. The subgroup of children marked as having a high-risk-of-bias were diagnosed postnatally in a medical evaluation following neurodevelopmental or behavioural concerns. fTCD data were not available on all of the children for the following reasons: inability to establish ultrasound signal (n = 18); poor signal (n = 9); child refusal (n = 19); child non-compliance (n = 2); task too difficult for the child (n = 8); insufficient time during testing (n = 7); error with the recording (n = 5). Please note that one child (a 47,XXX girl) is not included in the chart, since her early history was not known.

### Measures

Details of language and psychopathology assessments are reported elsewhere.

#### Handedness assessments

We used an adapted version of the Edinburgh Handedness Inventory (EHI)^20^ to measure hand preference, replacing one item (striking a match) with a more child-friendly one (dealing cards). The child was asked to demonstrate how they would perform ten actions. For each action, exclusive right hand use scored one point, left hand use zero, and both hands half. Totals out of ten were converted to indices between -100 (extreme left-handedness) and 100 (extreme right-handedness). Scores above zero indicated right-handedness.

The Quantification of Hand Preference (QHP) task^21^ measured strength of hand preference. This assesses tendency to reach across the midline with the preferred hand. 21 picture cards were arranged in front of the child, three stacked in seven positions at 30-degree intervals. The child placed named cards into a central box one at a time in the same quasi-random order used for all participants. The child was not told that the task assessed handedness. Each right-handed reach scored one point. Totals out of 21 were converted to indices, as above. Scores above ten indicated right-handedness.

#### Language laterality assessment

Language laterality was assessed using functional transcranial Doppler ultrasound (fTCD). Please see our related paper for a full description of the procedure and data analysis^19^. Briefly, ultrasound probes were positioned to detect blood flow in the left and right middle cerebral arteries (MCAs). The child performed an animation description task comprising a maximum of 30 trials. Each trial involved watching a 12 s cartoon, before describing it during a 10 s talk phase. Trials were excluded where the child spoke during a silent period, or said nothing during the talk phase.

Analysis of the fTCD data involved comparing blood flow during the period-of-interest where the child speaks with the baseline period spent watching the cartoon. Following initial data-processing, including exclusion of trials with poor signal, a laterality index (LI) was calculated. A 2 s window was constructed around the maximum difference in velocity between the two MCAs during the period-of-interest. The mean velocity in the left MCA during this window minus the mean in the right MCA gave the LI. Children with LIs based on fewer than 12 trials were excluded from analysis, since such LIs are unlikely to be reliable.

Participants were also assigned a laterality category. 95% confidence intervals were calculated around the LI. If the CIs did not cross zero, laterality was classified as left or right depending on direction, and bilateral where the CIs crossed zero. Bilateral laterality may result if data are merely noisy. (95% CIs were calculated using the standard error derived from trial-by-trial LIs computed for every valid trial during the same 2 s window as the overall LI.)

LIs were derived separately from odd and even trials to allow computation of split half reliability, and the mean number of words spoken by the child during valid trials was recorded.

#### Nonverbal ability and language status

Nonverbal ability was estimated using the two nonverbal subtests of the Wechsler Abbreviated Scale of Intelligence (WASI): Block Design and Matrices^22^. Scores were converted to Performance IQ.

A battery of 13 tests assessed language abilities^19^. In determining language status, performance at least 1 SD below the normative mean on two or more language tests indicated language problems. Otherwise, the child had typical language.

### Procedure

Ethical approval was obtained in 2011 from the Berkshire NHS Research Ethics Committee (reference 11/SC/0096). Data were collected between August 2011 and October 2016 from families throughout the UK. Families were first interviewed by telephone, and if inclusion criteria were met, an assessment was arranged at home or school. Written consent was obtained from a parent/caregiver, and children signed a simplified assent form. Eight research assistants and the senior author conducted assessments, scheduled in a single session lasting 2-3 hours.

### Statistical analysis

See the Appendix for details of (1) software used in analysis, and (2) data storage and availability.

For our main analysis, we tested the hypothesis that the SCT group would show a reduced bias for left-lateralised blood flow during a language task. Response variables were quantitative laterality (LI) and laterality category. Multiple regression was used to test for a Group effect (SCT or comparison group) on LI, controlling for age and sex. A multinomial logistic regression tested whether Group (SCT or comparison) predicted laterality category, controlling for age and sex. This model estimated two logit equations, each comparing relative frequency of left-lateralised language to one atypical laterality (bilateral and right), and assigned predicted log-odds to each predictor. We also tested whether handedness was atypical in the SCT children. As explained in our previous paper^19^, the two quantitative handedness measures are best modelled using inflated beta regression. We ran one model per measure; the logit function of the handedness measure was response variable and Group (SCT or twin), Age and Sex were predictors.

## Results

Table 1 shows summary statistics.

**Table 1.** All continuous variables are reported as means (SDs) and categoricvariables are reported with frequencies and percentages.

|  | SCT group | Comparison group |
| --- | --- | --- |
| **Sample Characteristics** |  |  |
| N, *children* | 75<sup>†</sup> | 132<sup>‡</sup> |
| Age, *years; months* | 11; 4 ($\pm 3$; 10) | 9; 1 ($\pm 1$; 7) |
| Sex, *n male* | 49 (65.33%) | 69 (52.27%) |
| Performance IQ | 93.16 ($\pm 16.17$) | 103.15 ($\pm 13.57$) |
| Language Status, *n with language problems* | 67 (89.33%) | 48 (36.36%) |
| **fTCD Language Laterality** |  |  |
| N trials completed | 24.75 ($\pm 5.25$) | 26.83 ($\pm 3.22$) |
| N words produced per trial | 17.70 ($\pm 6.19$) | 19.90 ($\pm 5.12$) |
| Laterality index | 1.87 ($\pm 3.04$) | 1.50 ($\pm 2.93$) |
| Left language, *n children* | 47 (62.67%) | 85 (64.39%) |
| Bilateral language, *n children* | 14 (18.67%) | 21 (15.91%) |
| Right language, *n children* | 14 (18.67%) | 26 (19.70%) |
| **Handedness** |  |  |
| Edinburgh Handedness Inventory (EHI) | 63.07 ($\pm 58.01$) | 62.50 ($\pm 61.04$) |
| n children right-handed by EHI | 64 (85.33%) | 110 (83.33%) |
| Quantified Hand Preference (QHP) | 13.19 ($\pm 6.96$) | 14.18 ($\pm 7.38$) |
| n children right-handed by QHP | 51 (68.00%) | 95 (71.97%) |

<sup>†</sup> Performance IQ only available for 68 of the 75 SCT children.
<sup>‡</sup> Performance IQ only available for 131 of the 132 comparison children.

### Preliminary Analysis of fTCD data

We considered whether there were any group differences that may affect interpretation of any laterality findings. Welch’s t-test revealed that fTCD data for the SCT children included fewer valid trials, *t* (106.19) = −3.12, *p* = .002, Cohen’s *d* = 0.45. The average child with an SCT also produced slightly fewer words per valid trial, *t* (126.89) = −2.59, *p* = .011, Cohen’s *d* = 0.38. Next we checked whether task performance (operationalised as N words produced per trial) correlated with LI; it did not, *r* = .002, *p* = .973. Mean number of words produced by left-lateralised children was 19.38 [SD = 5.63], compared to 17.71 [SD = 5.81] for those with bilateral language and 19.49 [SD = 5.31] for the right-lateralised children.

Finally, split half-reliability of the LIs was high in both groups, SCT *r* = .87, comparison *r* = .84, indicating that LIs represented stable cerebrovascular responses.

### Hypothesis-testing

#### Language laterality

See Figure 2 for a plot showing the time course of MCA blood flow, and Figure 3 for a pirate plot showing LI distributions.

**Figure 2.**
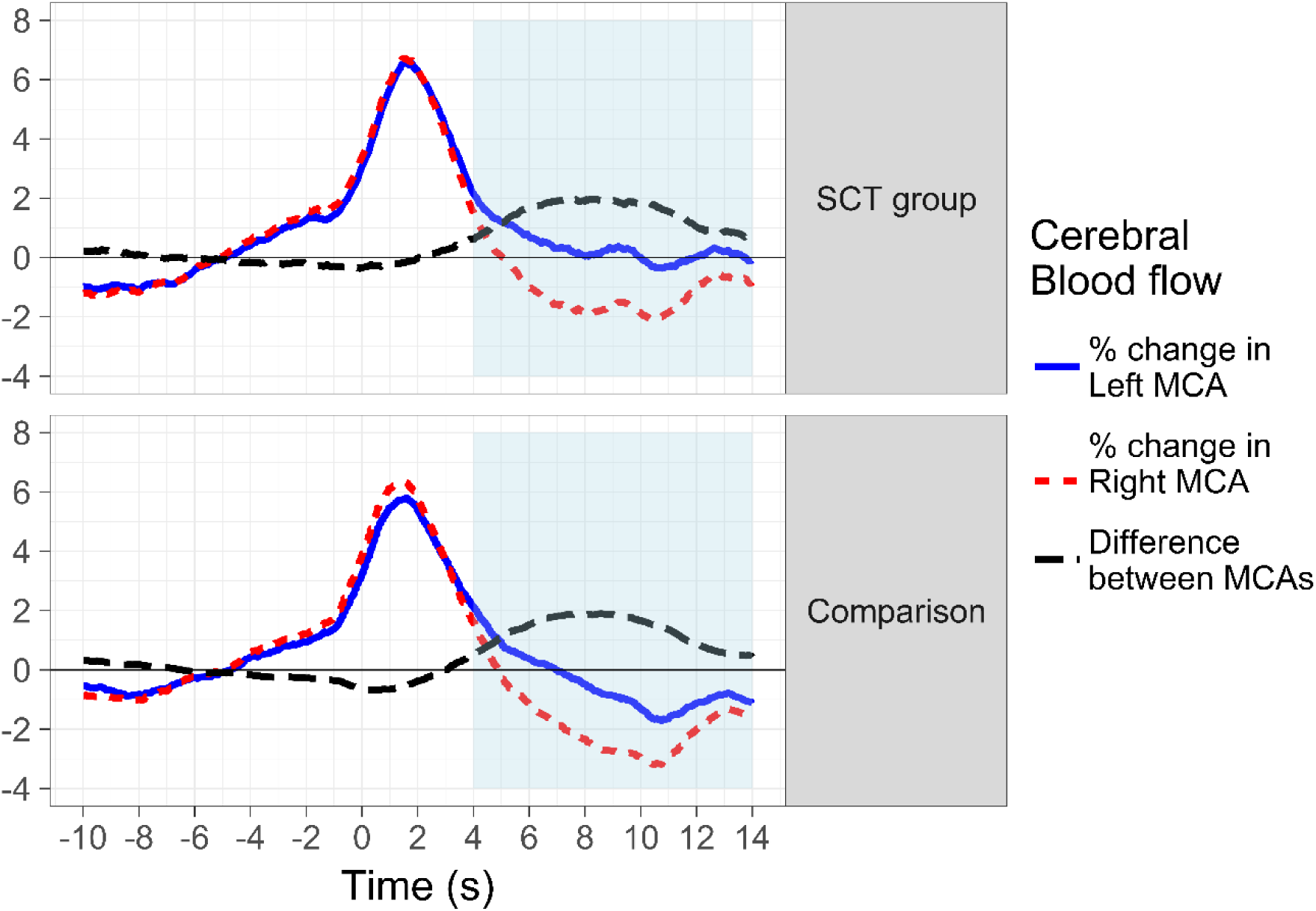
Plot showing grand average curves in the two groups (SCT and Comparison) for blood flow in the left and right MCAs during the time course of a trial.

**Figure 3.**
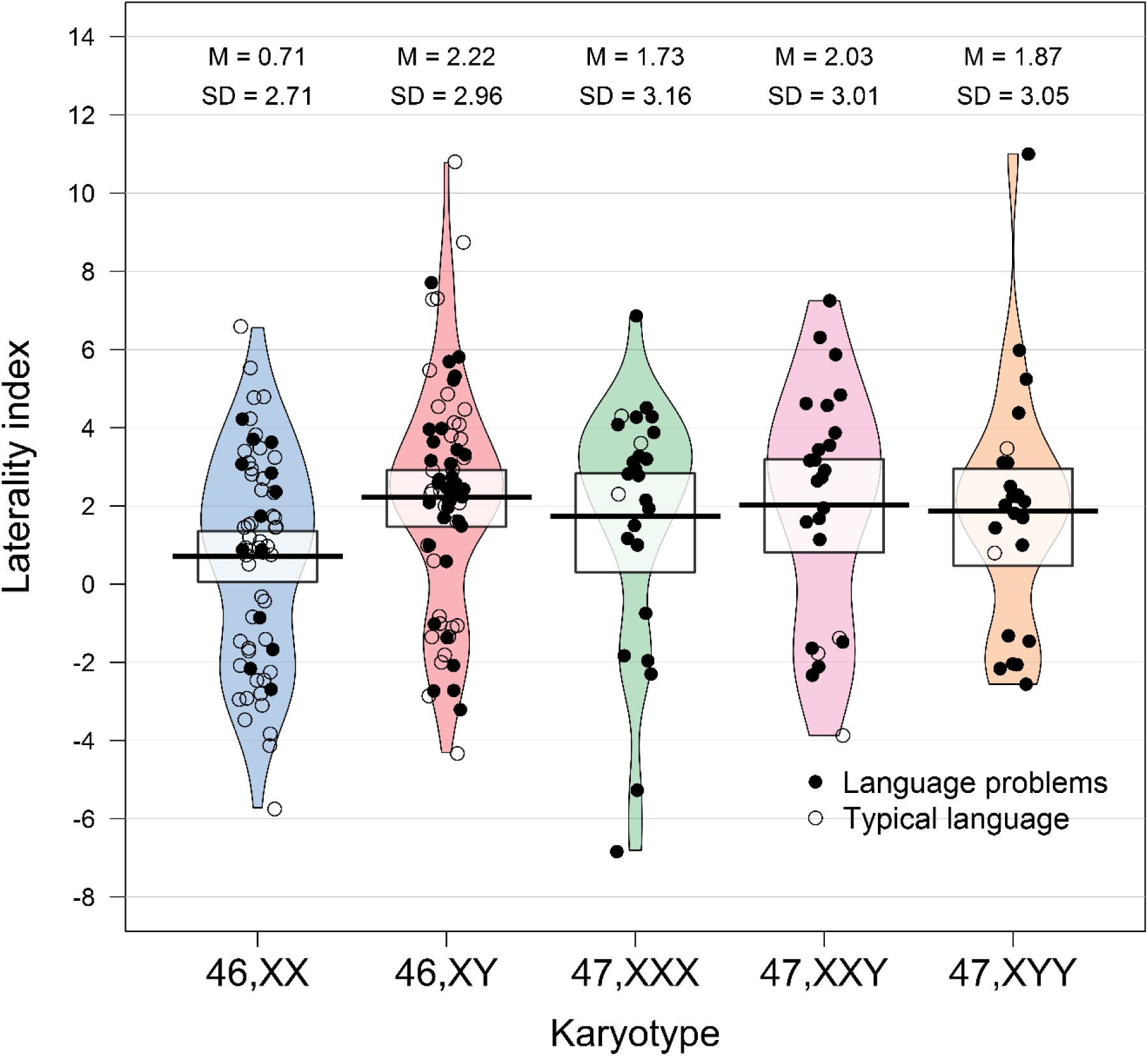
Pirate Plot showing distributions of laterality indices for the SCT and comparison children, split by karyotype. The 46,XX and 46,XY children constitute the comparison group. All data are shown with smoothed densities indicating the distributions in each subgroup. The central tendency is the mean and the intervals are Bayesian 95% Highest Density Intervals.

We tested our main hypothesis that mean LI would be reduced in the children with an SCT, when controlling for Age and Sex, using multiple regression. The Group factor was not significant, β = 0.07, SE = 0.16, *p* = .663, indicating, contrary to hypothesis, that both groups showed no difference in the usual left-sided bias for language. Age was not significant, β = 0.01, SE = 0.07, *p* = .860, though Sex was, β = -0.36, SE = 0.14, *p* = .011. The sex effect had already been reported for the twins comparison group^19^, so we ran a post-hoc t-test to test for a sex difference in the SCT group, which found no difference, *t* (48.78) = 0.29, *p* = .773.

Visual inspection of the pirate plot indicates no differences in LI by trisomy (47,XXX, 47, XXY or 47,XYY). Likewise, there were no laterality differences when analysing the SCT children by possible risk of bias based on circumstances of diagnosis. Mean LI of the high-risk-of-bias subgroup was 2.32 [SD = 3.31], compared to 1.72 [SD = 2.66] for the low-risk-of-bias children, *t* (55.83) = 0.83, *p* = .409.

We also checked for group differences in proportions of children with each laterality category. A multinomial logistic regression tested whether the odds of being atypically lateralised differed by Group, when controlling for Age and Sex. Comparing bilateral against left laterality, the predicted odds [95% CIs] for all factors were non-significant: Group 1.58 [0.70, 3.58], *p* = .268, Age 0.74 [0.49, 1.11], *p* = .144, and Sex 1.83 [0.85, 3.94], *p* = .124. Comparing right against left laterality, all factors were again non-significant: 0.79 [0.33, 1.91], *p* = .597, Age 1.30 [0.87, 1.95], *p* = .124, and Sex 1.52 [0.74, 3.13], *p* = .124. This analysis confirms that language laterality was not unusual in the children with an SCT.

#### Handedness

Inflated beta regressions indicated that there were no significant effects of Group, Age or Sex on either of the two handedness measures. Predicted odds [95% CIs] that an SCT child in relation to a comparison child was fully right rather than left-handed on the adapted EHI was 1.12 [0.74, 1.71], *p* = .597. Predicted odds [95% CIs] that an SCT child in relation to a comparison child was fully right rather than left-handed on the QHP was 0.83 [0.57, 1.22], *p* = .341.

## Discussion

The present study found no differences in cerebral lateralisation for language in 75 children with a sex chromosome trisomy. Previous research has offered weak support for structural differences in brain asymmetry in individuals with an SCT, though two small functional studies did report atypical lateralisation in 47,XXY individuals^4,5^. One was a resting state study^4^, and conceivably, differences in participant behaviour in the scanner contributed to the reduced functional asymmetry in the clinical group. The other functional neuroimaging study^5^ may have been distorted by an outlier, as discussed in the introduction. The present results question the leading theoretical accounts of the brain basis of the language phenotype in SCTs, and we outline points of contention below.

Rezaie and colleagues summarised the view that the SCT phenotype represents a left hemisphere dysfunction, since impairment in a typically left-lateralised function, language, is a hallmark feature^12^. These researchers contrasted SCTs with Turner’s Syndrome (where a female has one X chromosome rather than two), which is characterised by impaired visuospatial cognition, a typically right-lateralised function. This link between sex chromosome aneuploidies and lateralised cognitive functions has been taken as evidence that genes on the sex chromosomes influence lateralisation^3^. High dosage of such genes, when an individual has an extra sex chromosome, may disrupt lateralisation of left hemisphere function (whereas, for some unspecified reason, low dosage in Turner’s Syndrome affects the right hemisphere). However, the lack of effects of an extra sex chromosome on laterality in the present data provides no evidence for these hypothetical laterality genes.

An alternative theory is based on the hypothesis that the left hemisphere develops more rapidly, and has an inhibitory effect on the right, in most people^17^. This typical trajectory may be disrupted in individuals with an SCT due to a loss of the left-on-right inhibitory mechanism, such that the right hemisphere assumes some language function, contributing to reduced processing efficiency. Inefficient organisation of language function across the cerebral hemispheres has been theorised elsewhere as a factor in Developmental Language Disorder (DLD)^23^. However, this theory was not supported by analysis of a large sample of typically-developing and language-impaired children (n = 267), who showed no laterality differences^19^. As for the present study, 90% of the SCT children had language difficulties diagnosable as DLD, but there was no evidence of atypical laterality. Hemispheric specialisation for language was usually left-lateralised.

Theoretical accounts also implicate atypical cerebral laterality in the elevated psychiatric risk associated with SCTs. The most documented risk is for autism^24^. With relevance to the present study, theories have linked the communication difficulties in autism with atypical lateralisation^25^ and left hemisphere dysfunction^26^. In addition, the 47,XXX and 47,XXY karyotypes have been explored as genetic models for schizophrenia^5^, although an increased risk for schizophrenia is not well established^27^. Nonetheless, the relationship between atypical lateralisation and schizophrenia^28^, and the possibility that genes on the sex chromosomes influence lateralisation and psychosis^3^, make a putative link between SCTs, schizophrenia and lateralisation of substantial psychiatric interest. However, our null findings speak against the view that atypical laterality may be an endophenotype representing increased psychiatric risk in SCTs.

The main limitation in the present study is that fTCD may fail to detect fine-grained laterality differences, because it measures blood flow changes in the middle cerebral artery, which covers a wide territory. The possibility remains that small regional differences in lateralization would go undetected by this method. Note, however, that fTCD is sensitive to language-related activity, so should be able to detect differences at a network level^19^.

## Conclusion

In this relatively large study of children with sex chromosome trisomies, we found no evidence for atypical lateralisation for language. The proportion of children showing left, bilateral and right-lateralised language was identical with a comparison group, and mean laterality indices did not vary by karyotype. Our results disconfirm a leading hypothesis that atypical lateralisation for language is the neurobiological basis for the cognitive phenotype in sex chromosome trisomies.

## Acknowledgements

We offer warmest thanks to the families who took part in the study, and school staff who helped facilitate assessment arrangements. The study would not have been possible without the hard work and dedication of a series of research assistants who conducted the assessments, often travelling all over the UK to do so: Eleanor Payne, Nicola Gratton, Georgina Holt, Annie Brookman, Elaine Gray, Louise Atkins, Holly Thornton and Sarah Morris. We also thank Paul A. Thompson for expert advice on statistical analysis. This work was funded by Wellcome Trust Programme Grants no 082498/Z/07/Z and 082498/Z/07/C. The funder was not involved in study design, data collection, data analysis, manuscript preparation or publication decisions.

## Appendix

Study data were analysed using R software^1^, with the main database managed using REDCap, hosted at the University of Oxford^2^. Original data are available on *Open Science Framework* at https://osf.io/2w6u5/.

The multinomial logistic regressions were run in R using the nnet package^3^. The inflated beta regressions were implemented with the GAMLSS package^4^. The flow chart was produced using the DiagrammeR package^5^, the plot showing changes in MCA blood flow with ggplot2^6^, and the pirate plot with yarrr^7^. The R packages tidyverse^8^, stringr^9^, psych^10^, and xlsx^11^ were also used in analysis.

